# Population variability in X-chromosome inactivation across 9 mammalian species

**DOI:** 10.1101/2023.10.17.562732

**Authors:** Jonathan M. Werner, John Hover, Jesse Gillis

## Abstract

One of the two X chromosomes in female mammals is epigenetically silenced in embryonic stem cells by X chromosome inactivation (XCI). This creates a mosaic of cells expressing either the maternal or the paternal X allele. The XCI ratio, the proportion of inactivated parental alleles, varies widely among individuals, representing the largest instance of epigenetic variability within mammalian populations. While various contributing factors to XCI variability are recognized, namely stochastic and/or genetic effects, their relative contributions are poorly understood. This is due in part to limited cross-species analysis, making it difficult to distinguish between generalizable or species-specific mechanisms for XCI ratio variability. To address this gap, we measured XCI ratios in nine mammalian species (9,143 individual samples), ranging from rodents to primates, and compared the strength of stochastic models or genetic factors for explaining XCI variability. Our results demonstrate the embryonic stochasticity of XCI is a general explanatory model for population XCI variability in mammals, while genetic factors play a minor role.

## Introduction

Every female mammalian embryo undergoes X-chromosome inactivation (XCI) as an essential step for successful development^1–3^. XCI evolved to balance the gene dosage between females with two X-chromosomes and males with one X-chromosome^4^. While the exact timing can vary across species^5^, XCI usually occurs during preimplantation embryonic development^6^. During this process, one of the two X-alleles in each female cell is independently, randomly, and permanently chosen for transcriptional silencing to match the single X-allele in male embryos^1,7–9^. The choice of silenced X-allele is inherited through cell divisions, propagating the random choice of allelic inactivation down each cell’s subsequent lineage. This produces whole-body mosaicism for allelic X-chromosome expression in each adult mammalian female, originating from very early embryonic development^10^.

In humans, both X-alleles are equally likely to be inactivated, but XCI ratios vary widely among adult females, from balanced to highly skewed^11,12^. XCI ratios affect the phenotypes of X-linked diseases, as they can either protect or expose individuals to disease variants^10,13^. The factors that influence XCI variability are mostly studied in mice and humans, and include stochasticity^12^ and genetics^14–16^, but their relative roles are controversial^17^. Cross-species comparisons of XCI variability stand to reveal general or species-specific mechanisms of XCI. For instance, genetic determinants of XCI are well-established in lab mice^18–20^, but not in humans^17,21,22^, where they are harder to identify and measure. Exploring XCI variability in other mammals presents the opportunity to test models of stochasticity or genetics in the context of evolution.

Considering first a stochastic model for XCI variability, each cell within an embryo at the time of XCI independently selects an X-allele to inactivate, resulting in ratios of allelic-inactivation across embryos varying purely by chance (Fig. 1A). Closely following Mary Lyon’s discovery of XCI in 1961^1^, it was recognized that the inherent embryonic stochasticity and permanence of XCI is the simplest explanation for the observed variability in XCI among adults and positions this adult variability as a window into embryonic events^23–27^. For example, flipping 10 coins is more likely to result in 8 heads than flipping 100 coins is likely to result in 80 heads, meaning that the variability in heads-to-tails ratios depends on the number of coins flipped. Similarly, the variability of XCI ratios in a population of female mammalian embryos is determined by the number of cells at the time of XCI (Fig. 1A). Since each cell inherits its allelic-inactivation from its ancestor, measuring XCI variability in adults can approximate embryonic XCI variability and help infer cell counts at the time of XCI or other early lineage decisions^25,28^ (Fig. 1D). Stochastic models have been used to estimate cell counts during embryonic events in human and mice populations for decades^20,23,25,27–29^ – but their applicability has not been tested in other mammalian species.

**Figure 1:**
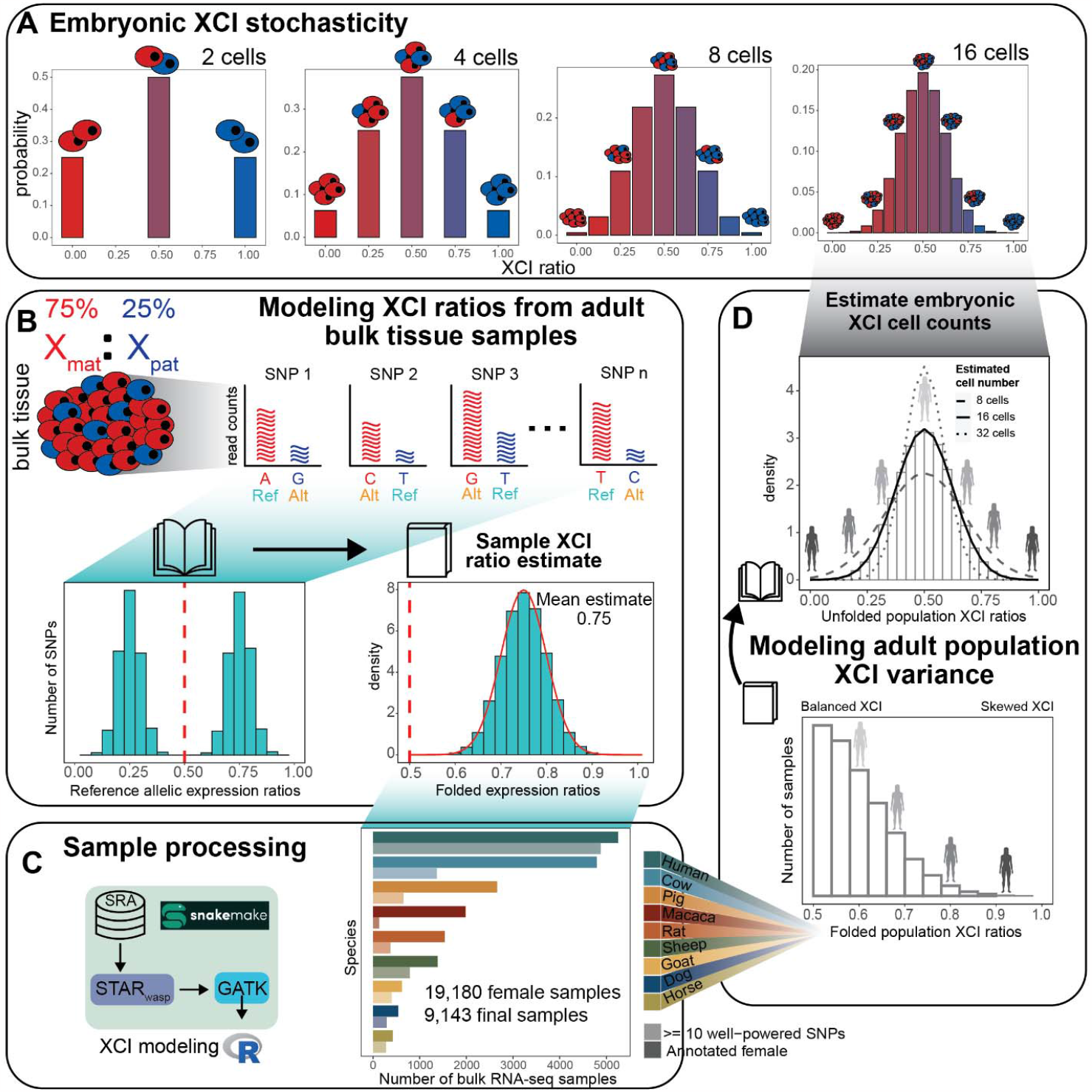
Reference aligned RNA-sequencing data enables scalable modeling of XCI ratios. **A** Schematic demonstrating the relationship between the number of cells present at the time of XCI and the probability of all possible XCI ratios. Increased cell numbers result in decreased XCI ratio variance. **B** Schematic for modeling XCI ratios from bulk reference-aligned RNA-seq data. The reference SNPs will contain both maternal and paternal SNPs, representing allelic expression from both parental haplotypes. Folded normal models are fit to the folded reference allelic expression ratios (like folding a book closed), with the mean of the maximum-likelihood distribution as the sample XCI ratio estimate. **C** Schematic for sample processing (genome alignment and variant identification) and a bar graph depicting the number of annotated female samples initially downloaded for each species (bold color), with the number of samples per species with at least 10 well-powered SNPs for XCI ratio modeling after processing (faded color). **D** Schematic demonstrating the population modeling of XCI variability. Folded population distributions are first produced per species and then are unfolded. Normal distributions are fit to the unfolded population distribution to estimate the number of embryonic cells required to produce the observed variance.

In addition to stochasticity, genetic effects can influence the choice of allelic inactivation and contribute to population variability in XCI ratios. Allelic inactivation during XCI is mediated by the cis-acting long non-coding RNA XIST^30^, which silences its corresponding X-allele through epigenetic modifications^31,32^. Heterozygous variants affecting XIST expression can bias allelic inactivation^15^. For example, inbred mice show preferential inactivation of specific X-alleles depending on the parental strains and their corresponding X-chromosome controlling element (XCE) allele^18,20,33^. In humans, genetic influence on XCI is mostly observed in small family studies or disease cases, with no strong evidence for the broad allelic effects seen in mice^21,22^. Another genetic influence on XCI is allelic selection, where natural or disease-causing variants favor certain X-alleles^14,16,34–38^. However, evidence for allelic selection through natural variation remains elusive in human populations. Thus, the relative contributions of stochasticity and genetics to population XCI variability in mammals remain unclear with currently limited data from mouse and human studies.

In this study, we assess population scale XCI variability and its determinants across nine mammalian species. We source female annotated bulk RNA-sequencing samples from the Sequencing Read Archive (SRA), resulting in a total of 19,180 initial samples (Fig. 1C), including human samples from the GTEx^39^ dataset. Our approach leverages natural genetic variation to sample X-linked heterozygosity and eliminates the requirement for costly phased or strain specific genetic information to assess XCI ratios across diverse mammals at population scale. We start by establishing the population-level XCI ratio distributions for all nine mammalian species and use models of embryonic stochasticity to predict the number of cells fated for embryonic lineages (Fig. 1D, Fig. 2). We then investigate how broad genetic diversity, as indicated by measures of inbreeding (Fig. 3), as well as specific individual variants (Fig. 4), may impact population XCI variability. Overall, our analyses explore how both models of stochasticity and genetic factors can explain population XCI variability across diverse mammalian species.

**Figure 2:**
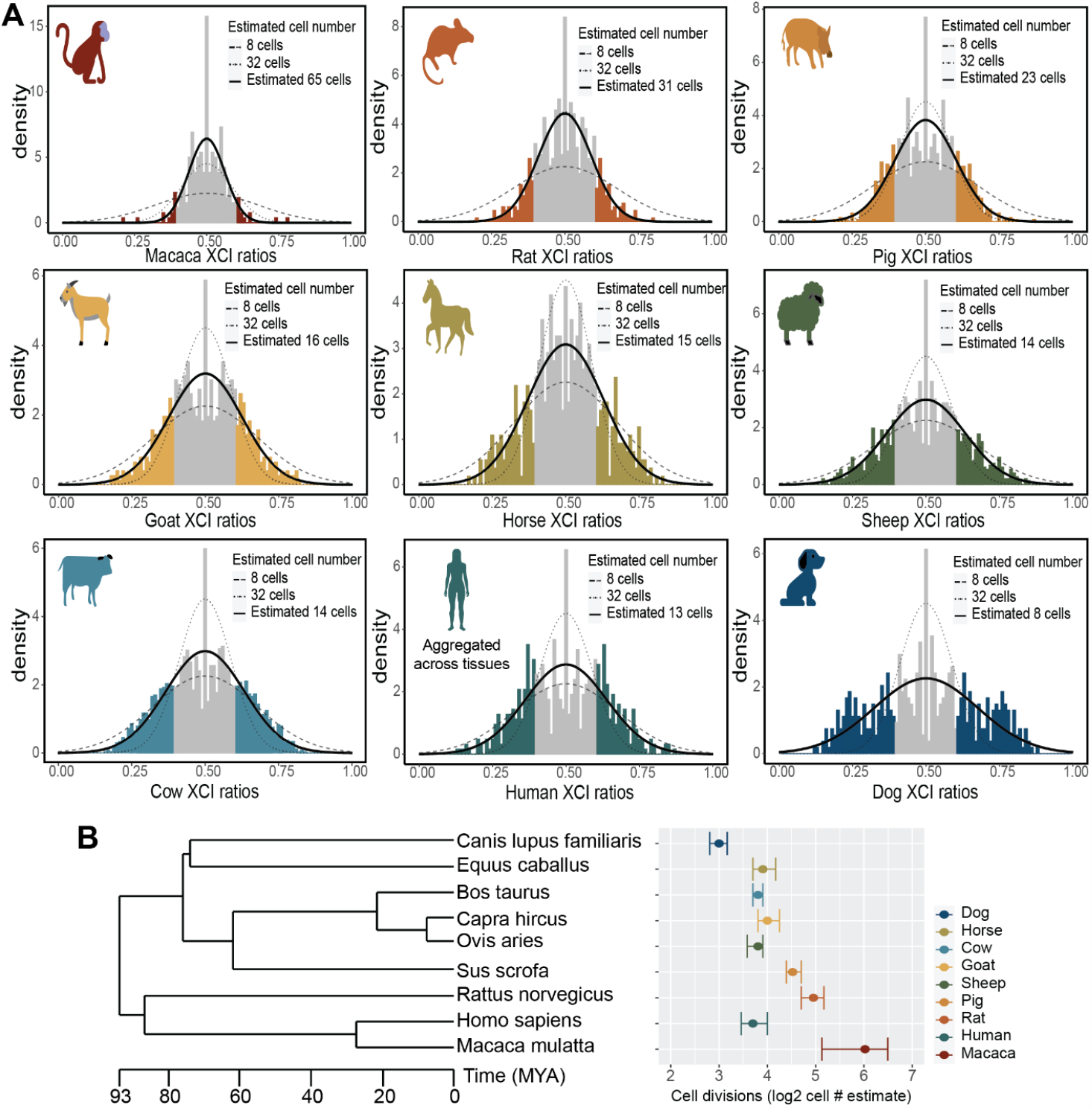
Models of embryonic stochasticity explain adult population XCI variability. **A** Unfolded distributions of XCI ratios per species, with the maximum-likelihood normal distribution depicted in bold, fitted to the tails of the distributions (shaded in sections of the distributions). **B** Phylogenetic tree of the sampled mammalian species with their estimated embryonic cell counts on a log-2 scale, depicting the number of cell divisions that separate the estimated cell counts between the species. Error bars are 95% confidence intervals around the cell number estimate.

**Figure 3:**
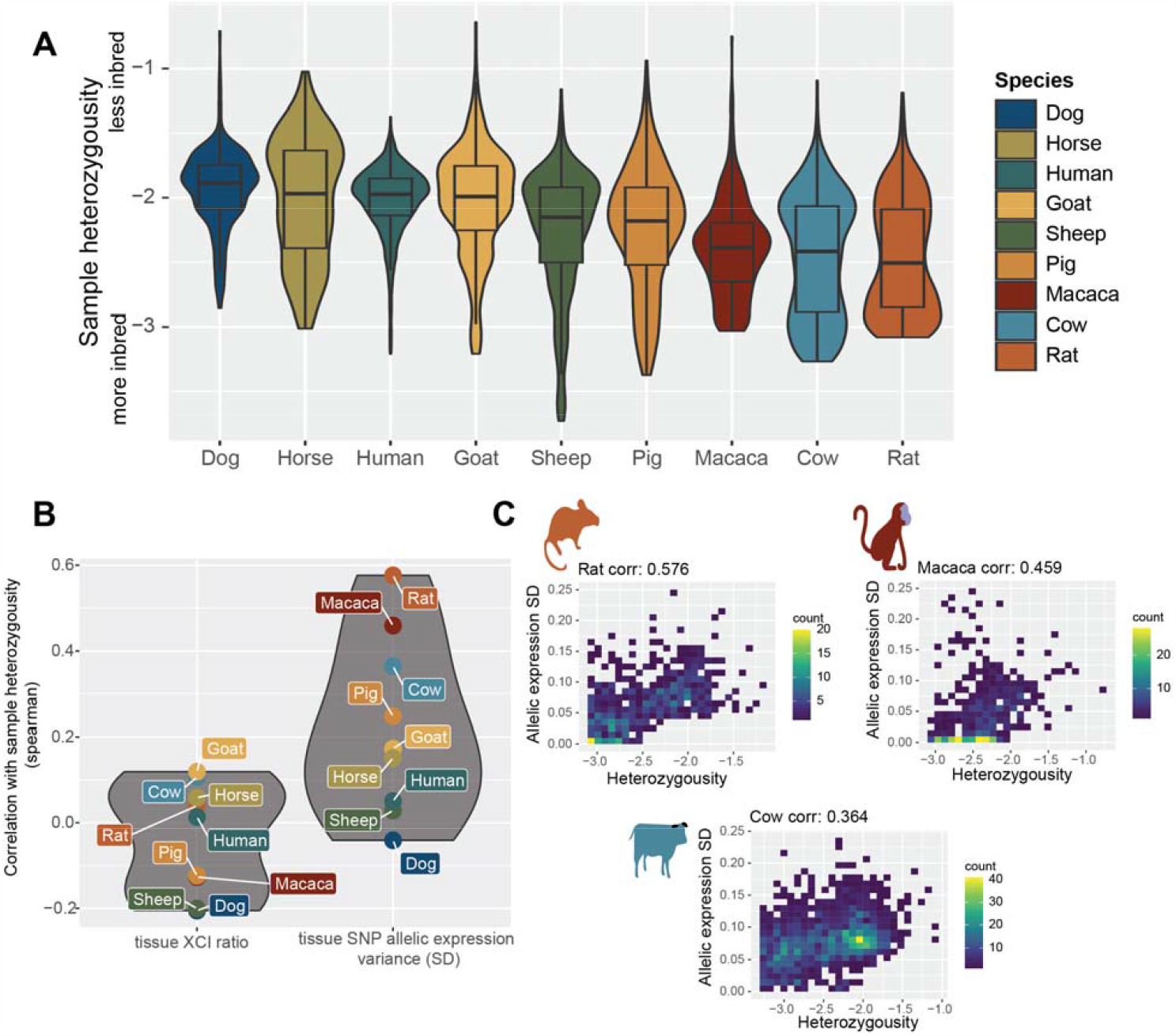
XCI ratios are not associated with X-linked heterozygosity. **A** Distributions of sample X-linked heterozygosity per species ordered by the median value. The y-axis is in log-10 scale, depicting the ratio of SNPs per sample to all unique identified SNPs per species. Boxplots depict the distributions’ quartiles. **B** The spearman correlation coefficients between sample X-linked heterozygosity and either the estimated standard deviation (SD) in X-linked allelic expression or the estimated XCI ratio of the sample (the SD and mean of the maximum-likelihood folded-normal model per sample). **C** 2D Scatter plots of sample heterozygosity compared to the sample estimated X-linked allelic expression SD for the three species with moderate correlation coefficients. Color bars represent the number of samples in each 2D bin. Plots for the other species are in Supp. Fig. 6.

**Figure 4.**
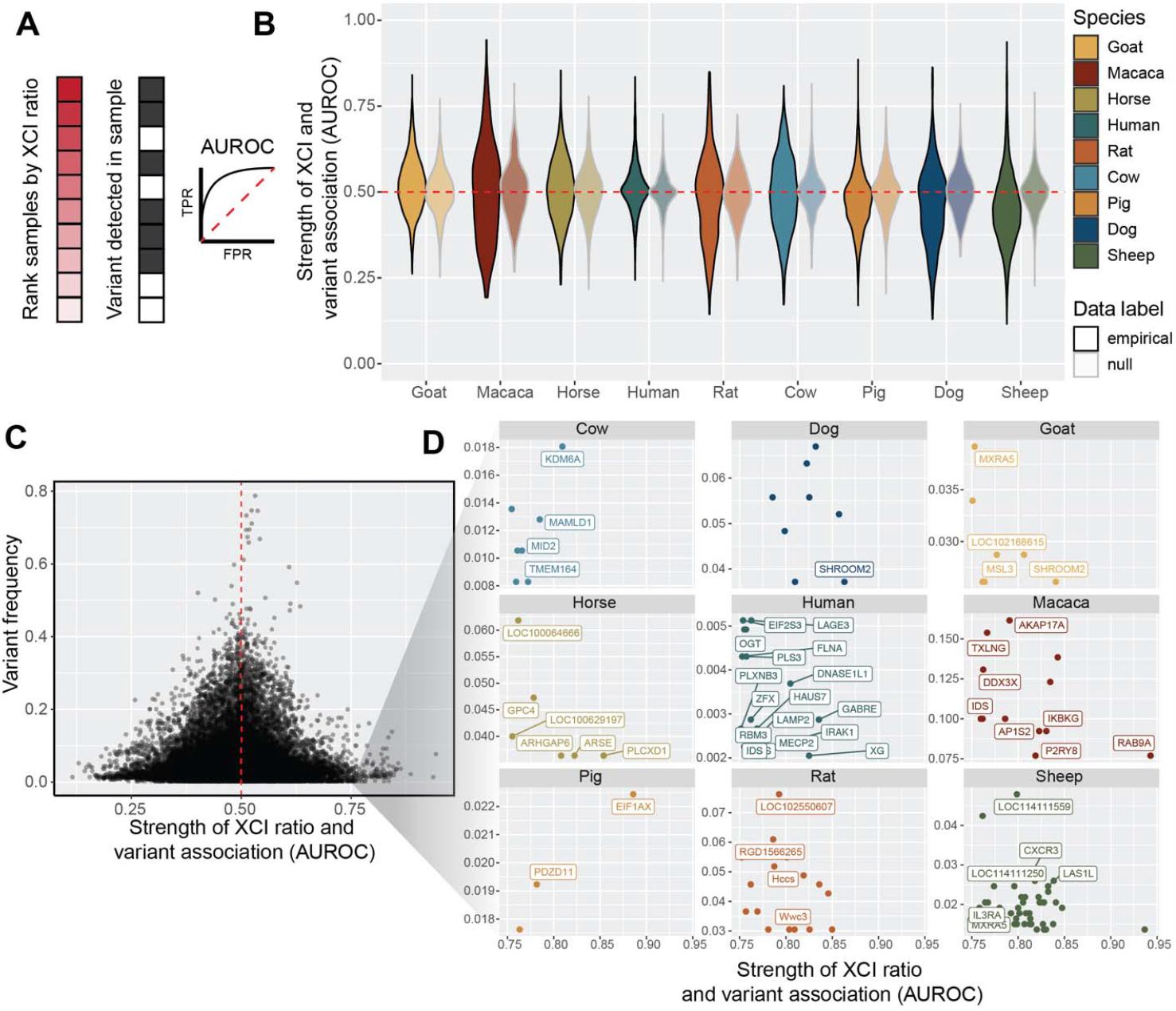
Low frequency variants exhibit moderate associations with XCI ratios. **A** Schematic depicting the AUROC quantification for testing the association between individual variants and extreme XCI ratios. Samples are ranked by their estimated XCI ratio, with the dark shaded red squares representing samples with more extreme XCI ratios. The position of samples with a given individual variant (grey squares) within the ranked list is used to compute the AUROC statistic. A variant with an AUROC value of 1 means all samples with that variant were at the top of the ranked list, whereas an AUROC value of 0.5 represents a random ordering of samples within the ranked list. **B** Distributions of variant AUROCs for each species compared to a species-specific null distribution of AUROC values (faded distributions, see methods), ordered by the mean value of the empirical distributions. The red dotted line depicts an AUROC of 0.50, performance due to random chance. **C** Scatter plot of variant AUROCs compared to each variant’s prevalence (percent of samples with that variant, relative for each species) for all variants across all species. The red dotted line depicts an AUROC of 0.50, performance due to random chance. A threshold of AUROC >= 0.75 was used to identify SNPs with moderate associations with XCI ratios. **D** Scatter plots depicting the same information as in C for the variants with moderate associations with XCI ratios, but split by each species and including gene annotations. SNPs not within annotated genes are unlabeled. Gene labels not present due to overlapping labels are Macaca: ZBED1, Sheep: LOC101108113, LOC101115509, LOC101117055, LOC105605313, LOC121818231, PPP2R3B, PRKX)

## Results

### Reference aligned RNA-sequencing data enables scalable modeling of XCI ratios

We use bulk RNA-sequencing (RNA-seq) data to measure the X-linked allelic expression of a sampled tissue by computing allele-specific expression ratios of heterozygous single nucleotide polymorphisms (SNPs). The parental proportion of X-linked allelic reads are expected to follow a binomial distribution dependent on the number of sampled reads and the XCI ratio of the tissue (see methods). The binomial distribution is an appropriate model when the parental identity of sequencing reads is known, which is not the case when aligning to a reference genome. A reference genome will contain SNPs from both parents, making the parental identity of aligned reads ambiguous and producing reference allelic expression ratios that represent expression of both parental X-alleles (Fig. 1B).

We fold the distribution of reference allelic-expression ratios around 0.50 to aggregate data across both alleles and enable a robust estimate of the XCI ratio magnitude for the bulk RNA-seq sample (Fig. 1B). We fit folded-normal distributions to the reference allelic expression ratios of multiple SNPs per sample, which serves as a continuous approximation of the underlying depth-dependent mixture of folded-binomial distributions per SNP. The mean of the fitted distribution is the estimate of the XCI ratio (Fig. 1B). We also incorporate specific steps to address confounding factors that can impact X-linked allelic expression, including reference bias and escape from XCI^40,41^ (Supp. Figs. 1-2, see methods). Interestingly, we find the strongest signals of escape from XCI near chromosomal ends across all species (Supp. Fig. 2), suggesting escape within pseudo-autosomal regions is conserved across mammals^40,42^. Previously, we validated our SNP filtering and XCI modeling approach using phased RNA-seq data (where haplotype information is known for each variant) from the EN-TEx consortium^43^, achieving nearly perfect agreement in XCI ratio estimates for samples with folded XCI ratios of 0.60 or higher, demonstrating the accuracy of our approach.

By calling SNPs from RNA-seq reads and employing folded distributions to model reference-aligned allelic expression, we can estimate the magnitude of XCI in any female mammalian bulk RNA-seq sample. We source female annotated bulk RNA-seq samples of 8 non-human mammalian species from the SRA database (Fig. 1C), additionally including cross-tissue human samples from the GTEx dataset. After processing, the number of samples with a minimum of 10 well-powered SNPs for estimating XCI ratios are 130 macaca (mean of 28 SNPs +- 17 SD), 275 horse (mean of 54 SNPs +- 36 SD), 269 dog (mean of 29 SNPs +- 13 SD), 328 rat (mean of 26 SNPs +- 13 SD), 383 goat (mean of 34 SNPs +- 14 SD), 624 pig (mean of 50 SNPs +- 28 SD), 731 sheep (mean of 79 SNPs +- 42 SD), 1328 cow (mean of 32 SNPs +- 19 SD), and 4877 human (mean of 56 SNPs +- 23 SD, 314 total individuals) samples (Fig. 1C, Supp. Fig. 1). Aggregating reference allelic expression ratios for samples with similar estimated XCI ratios (0.05 bins) clearly reveals the expected haplotype expression distributions, demonstrating the applicability of folded models (Supp. Fig. 3). Following XCI ratio modeling, we then generate population-level distributions by unfolding the distribution of folded XCI ratio sample estimates per species (Fig. 1D).

To ensure the allelic variability we report from X-linked SNPs is specific to XCI, we estimate autosomal allelic imbalances for all samples using the same pipeline and approach as for the X-chromosome analysis (Supp. Fig. 4, see methods). Comparing allelic imbalances across the two autosomes closest in size to the X-chromosome reveals the vast majority of samples across all species are biallelically balanced for autosomal expression, as expected (Supp. Fig. 4). Several species (Pig, Cow, Goat, Rat, Sheep, and Dog) exhibit small subsets of samples that are consistently imbalanced across the two autosomes and the X-chromosome, indicative of a global influence on allelic-expression independent of XCI (Supp. Fig. 4). These samples with global allelic imbalances are excluded from all downstream analysis, ensuring the population distributions of XCI ratios reflect variability specific to XCI.

### Models of embryonic stochasticity explain adult population XCI variability

After generating population distributions of XCI ratios for the 9 mammalian species, we next explore how well models of embryonic stochasticity explain the observed adult XCI ratio variability. The initial variability in XCI ratios among mammalian embryos is dependent on the number of cells present during XCI (Fig. 1A), where adult variability can be modeled to infer embryonic cell counts.

An important consideration when estimating embryonic cell counts from XCI variability in adult tissues is the fact adult tissues only represent the embryonic lineage of the blastocyst as opposed to extra-embryonic lineages. This positions XCI variability of adult tissue samples as informative for the number of cells present within the last common lineage decision for all adult cells, i.e. the number of cells present within the epiblast of the mammalian blastocyst. If XCI occurs after epiblast specification, the variability in XCI ratios is determined by the number of epiblast cells at the time of XCI. On the other hand, if XCI occurs before epiblast specification, XCI variability within the embryonic lineage is influenced by both the initial stochasticity of XCI and the stochasticity associated with cell sampling during epiblast lineage specification. The temporal ordering of XCI among these lineage events cannot be resolved without cross-tissue sampling of both the extra-embryonic and embryonic tissues. As such, estimating cell counts solely on XCI variability in adult tissues provides an estimate of the number of cells present within the epiblast of the embryo.

Figure 2A presents the unfolded population distributions of XCI ratios in the 9 mammalian species we sampled, ranging from the least variable (macaca) to most variable (dog). We fit normal distributions as continuous approximations to the underlying binomial distribution that defines the relationship between cell counts and XCI ratio variability (Fig. 1A,D, see methods). We focus on the tails of the distributions, as our previous validation using phased data indicated increased uncertainty for folded XCI ratio estimates between 0.5-0.6, which translates to unfolded estimates between 0.4-0.6. At a broad level, population XCI ratio variability varies substantially across the sampled mammalian species. Our estimates for the number of epiblast cells present at the time of XCI include 65 (macaca), 31 (rat), 23 (pig), 16 (goat), 15 (horse), 14 (sheep), 14 (cow), 13 (human) and 8 (dog) cells, with associated 95% confidence intervals presented in figure 2B. The error between the empirical XCI ratio distributions and the normal fitted distributions is strikingly small, with a mean of 0.00538 (+-0.0101 SD) across the species (Supp. Fig. 5). This indicates models of embryonic stochasticity can explain observed XCI ratio variability in adult populations exceptionally well.

For the least and most variable species (macaca and dog), the estimated autosomal imbalances offer additional context for the reported XCI population variability. The reported X-linked variability in macaca is in excess to the reported autosomal allelic variability (Supp. Fig. 4). This demonstrates the X-linked population variability for macaca, while strikingly small, is specific to XCI and informative for estimating cell counts. On the other hand, the dog population is the only one that contains samples with strong allelic imbalances on only one autosome, where autosomal imbalances in all other species are global (Supp. Fig. 4). This is suggestive of broader genomic incompatibilities within the dog population. The reported X-linked population variability in dog is likely a combination of XCI and broader allelic incompatibilities, positioning our estimate of 8 cells as a likely underestimate due to excess variability outside of XCI.

Modeling XCI ratio variability across numerous species allows comparisons in light of evolution for determining generalizable or species-specific characteristics in XCI. Broadly, we demonstrate XCI ratios are variable in each species we assess, revealing variability in XCI ratios itself as a conserved characteristic of XCI. The exact variance in XCI ratios varies across the species, with differences in the timing of XCI and/or embryonic/extra-embryonic lineage specification (differences in cell counts) as one putative explanation. We compare our estimated cell counts to the evolutionary relationships among the species we assess (Fig. 2B), suggesting that variability in timing for these early embryonic events can be recent evolutionary adaptations. This is highlighted by the large differences in cell counts between macaca and humans. When viewed through the lens of cell divisions (log2 of the estimated cell counts, Fig. 2B), the differences in XCI ratio variability among the species can be explained by differences in a range of only 3 cell divisions, a narrow developmental window. This demonstrates even slight changes in the timing of XCI or embryonic/extra-embryonic lineage specification across mammalian species can produce large differences in population XCI ratio variability, as explained through the inherent stochasticity of XCI.

### XCI ratios are not associated with X-linked heterozygosity

After determining stochastic models can explain population XCI ratio variability across mammalian species, we turn to testing whether we can identify any genetic correlates with XCI ratios. Our approach leveraging natural genetic variation to quantify XCI ratios enables us to assess a large catalog of genetic variants for associations with XCI ratios across mammalian species (10,735 macaca SNPs, 12,024 rat SNPs, 23,603 pig SNPs, 16,123 goat SNPs, 10,281 horse SNPs, 53,505 sheep SNPs, 18,509 cow SNPs, 16,168 human SNPs, and 10,050 dog SNPs). One putative genetic contribution to XCI ratio variability is allelic selection during development, where increased X-linked heterozygosity (i.e., genetic distance), is more likely to produce selective pressures between the two X-alleles. It follows that samples with higher X-linked heterozygosity would be expected to exhibit more variability in XCI ratios.

We score X-linked heterozygosity per sample as the ratio of the detected SNPs within a sample to the number of unique SNPs identified across all samples, relative for each species (Fig. 3A). This quantification also serves as a measure of inbreeding, with decreased heterozygosity associated with a higher degree of inbreeding^44^. The trend in heterozygosity across species is as expected, with rats (likely laboratory strains) as the most inbred (Fig. 3A). Next, we examine the correlations between sample heterozygosity and the estimated XCI ratio, as well as the estimated XCI variability across SNPs in each sample (mean and standard deviation of the fitted folded-normal distribution per sample, Fig. 3B). Across all species, X-linked heterozygosity showed a near-zero correlation with the estimated XCI ratio, indicating a lack of association between X-linked genetic variability and XCI ratio variability (Fig. 3B). However, we observe moderate correlations between sample heterozygosity and the estimated variability in SNP allelic ratios in three species: rat (corr: 0.576), macaca (corr: 0.459), and cow (corr: 0.364), notably the most inbred species (Fig. 3A, Supp. Fig. 6). The increased variability in allelic expression present only within the most inbred species could potentially reflect gene-specific regulatory events between parental haplotypes^45^ rather than a direct genetic effect on XCI.

### Low frequency variants exhibit moderate associations with XCI ratios

After investigating relationships between genetic variation and XCI ratios at a broad level across the whole X-chromosome, we next asked if individual variants might be associated with extreme XCI ratios. Variants that affect the expression and/or function of the genetic elements that control XCI can result in highly skewed XCI ratios, as documented in human studies^15^. This can also occur in other X-linked genes, if the resulting differential in gene activity exerts a selective pressure across the X-alleles, as documented in disease cases^14,16^. We test the association between XCI ratios and individual variants for all variants detected in each species with a minimum of 10 samples, quantified through the area-under-the-receiver-operating-curve statistic (AUROC). For each species, we rank the samples based on their estimated XCI ratio and score the placement of samples carrying a given variant within the ordered list (Fig. 4A). If all the samples with that variant are at the top of the ordered list, the XCI ratio can be said to have perfectly predicted the presence of that variant, quantified with an AUROC of exactly 1. An AUROC of 0.50 indicates the XCI ratio performs no better than random chance for predicting the presence of the variant.

The distribution of AUROCs for each species show striking similarities to a null comparison (Fig. 4B, see methods), indicating a pervasive lack of association between XCI ratios and individual variants. However, a small subset of variants in each species exhibits moderate associations (AUROCs >= 0.75). By comparing each variant’s AUROC with its frequency in the species, we find that the variants with moderate associations occur at low frequencies within the sampled populations (Fig. 4C, Supp. Fig. 7). We investigate whether this relationship is simply due to a lack in power with bootstrap simulations, demonstrating moderate AUROCs (>= 0.75) are robust to their small sample sizes (Supp. Fig. 7). Figure 4D displays these variants along with their gene annotations for each species. Notably, several genes in humans with moderate AUROCs have prior evidence for associations with skewed XCI, namely MECP2^46^, IDS^47^ (also identified in macaca), IRAK1^48^, and FLNA^49^. This suggests our analysis is able to recover putative examples of selection impacting XCI ratios via disease-variants, though with small effect sizes and low frequencies in our sampled population. In general, we are unable to identify strong associations between genetic variation and XCI ratios across all 9 mammalian species, both along the whole X-chromosome and for individual variants.

## Discussion

We modeled tissue XCI ratios from bulk RNA-seq samples across 9 mammalian species and found population-level variation in XCI ratios, reflecting differences in developmental events such as XCI timing or lineage specification. We showed that embryonic stochasticity models fit the XCI data well and estimated epiblast cell counts at the time of XCI across species. We also searched for genetic factors influencing XCI ratios and found a pervasive lack of strong genetic associations with XCI ratios, indicating that XCI variability is mainly driven by stochasticity rather than genetic variation in mammals.

The lack of cross-mammalian comparisons of population XCI variability has previously limited our understanding on the sources of XCI variability in mammals. The existence of XCE-alleles in laboratory mice^18–20,33^ has supported the hypothesis that a similar genetic mechanism can exist in humans and drive population XCI variability^21^, though evidence for XCE-alleles in human populations remains inconclusive^22^ and data from other mammalian species is historically absent. Although genetic influences on XCI, particularly variants affecting XIST^15^ or disease-associated variants^34–37^, have been identified, they do not constitute a general mechanism that can fully account for observed population-level XCI variability. Comprehensive assessment of genetic influence on XCI would require combined DNA and RNA sequencing data, which is challenging to perform at a large scale across mammalian populations. Our approach for extracting heterozygous variants from RNA-seq data^28^, while providing a sample of genetic variability, is still able to assess hundreds of X-linked genes per species for associations with XCI and culminated in only weak evidence for limited genetic influence on XCI ratios. In contrast, we demonstrated models of embryonic stochasticity can explain population XCI variability with exceedingly small amounts of error consistently across mammalian species, providing a much more general explanation for population XCI variability.

Besides X-linked disorders and XIST-variants, other factors that may affect XCI ratio variability are genomic incompatibilities^45^ and stochastic allelic drift during development^20^. We found a link between the variance in X-linked allelic expression and the inbreeding level of some species (Fig. 2B), as well as autosome-specific allelic imbalances in dog (Supp. Fig. 4). This implies that X-linked allelic expression variability may result from both the bulk XCI ratio and the genomic incompatibilities between the parental genomes^45^, depending on the species. We controlled for global allelic imbalances by excluding samples that showed them (Supp. Fig. 4), which confirms that the allelic-expression variability on the X-chromosome is specific to XCI. Moreover, developmental allelic drift may introduce XCI ratio variability beyond the initial random choice of allelic inactivation^20^. While our previous cross-tissue analysis of XCI ratios in humans^28^ showed consistent XCI ratios across tissues, suggesting allelic drift is not a major factor in XCI ratio variability, similar data for non-human mammals is missing. These factors indicate that our cell count estimates are lower bound estimates for the number of cells needed to produce the observed XCI ratio variability as purely due to embryonic stochasticity.

A general model of X-linked genetic variability depletion (due to strong purifying selection in males^50–53^) accounts for the lack of evidence for broad allelic-selection or individual variants influencing XCI ratio variability in mammals, as both parental alleles are mostly equivalent. This does not apply to disease variants, but they cannot explain the widespread XCI ratio variability across mammalian species. We find genes associated with increased XCI ratios that have prior evidence for causing highly skewed XCI in disease cases, but their effect sizes and population frequencies are small in our samples. Therefore, the inherent stochasticity of XCI during embryogenesis is the main source of the observed XCI ratio variability in mammalian populations.

## Methods

### Snakemake pipeline for RNA-seq alignment and variant identification

All non-human mammalian fastq data was downloaded from the Sequencing Read Archive (SRA, https://www.ncbi.nlm.nih.gov/sra), where only samples annotated as female were selected, using the metadata provided through SRA. Details for download and processing of the GTEx^39^ data can be found here^28^. The entire sample processing pipeline uses a standard collection of bioinformatics software tools, all available for installation via Conda (STAR^54^ v2.7.9a, GATK^55^ v4.2.2.0, samtools^56^ v1.13, igvtools^57^ v2.5.3, and sra-tools 2.11.0). All Snakemake workflow rules, environment setup procedure, analysis commands and options, and underlying libraries are available on Github at https://github.com/gillislab/cross_mammal_xci, and https://github.com/gillislab/xskew. Briefly, a .fastq file acts as input, for either single- or pair-end sequencing experiments, and a .vcf and .wig file are produced as outputs for subsequent compiling of allele-specific read counts in R v4.3.0. The R script used for combining the .vcf and .wig information is also made available at https://github.com/gillislab/cross_mammal_xci/tree/main/R. Genome generation and alignment was performed with STAR, with the addition of the WASP^58^ algorithm for identifying and excluding reference biased reads. We extract chromosome-specific alignments from the .bam file (X chromosome or specific autosomes) and use GATK tools to identify heterozygous SNPs from that chromosome. The suite of GATK tools for identifying heterozygous variants from RNA-sequencing data was used following the <u>GATK Best Practices recommendations</u>. Specifically, the tools utilized include AddOrReplaceReadGroups -> MarkDuplicates -> SplitNCigarReads -> HaplotypeCaller -> SelectVariants -> VariantFiltration.

Reference genomes and gene annotations (.gtf files) for each species were sourced from the NCBI Refseq database (https://www.ncbi.nlm.nih.gov/refseq/). In each case the latest assembly version path was used, and the genomic.fna and genomic.gtf was downloaded. Annotated and indexed genomes were generated with STAR using --runMode genomeGenerate with default parameters.

### SNP filtering

Only SNPs with exactly two identified genotypes were included for analysis and indels were excluded. We required each SNP to have a minimum of 10 reads mapped to both alleles for a minimum read depth of 20 reads per SNP. Gene annotations for all SNPs were extracted from the species-specific .gtf files. For XCI ratio modeling, we only used SNPs found within annotated genes. For any sample with multiple SNPs identified in a gene, we took the SNP with the highest read count to be the max-powered representative of that gene, so each individual SNP is representative of a single gene. In addition to implementing the WASP algorithm for excluding reference biased reads, we filter out SNPs within each species whose mean expression ratios across samples deviate strongly from 0.50 (mean allelic ratio < 0.40 and > 0.60, Supp. Fig. 1). This SNP filtering also excludes potential eQTL effects that may impact allelic-expression outside of the underlying XCI ratio.

### Identifying and excluding chromosomal regions that escape XCI

We reasoned robust escape from XCI would produce more balanced biallelic expression in samples with skewed XCI. We performed an initial pass at XCI ratio modeling including all well-powered SNPs in a sample to identify samples with skewed XCI ratios (XCI ratios >= 0.70 for all species except rat and macaca, where a threshold of 0.60 was used due to a reduced incidence of skewed XCI in these species). Using the subset of skewed samples for each species, we averaged the folded allelic-expression ratios for all SNPs present in 1 mega-base (MB) bins across the X-chromosome (Supp. Fig. 2). Chromosomal-bins that displayed balanced allelic expression in opposition to the clearly skewed allelic expression of the rest of the chromosome were excluded from analysis. Specifically, chromosomal bins with an average allelic-expression < 0.65 for pig, goat, horse, sheep, and cow, < 0.60 in rat and macaca, and <0.675 in dog were excluded (Supp. Fig. 2) The ends of the X-chromosome in all species, except rat, demonstrated strong balanced biallelic expression, indicative of escape within putative pseudo-autosomal regions. We excluded any bin within these putative pseudo-autosomal regions regardless of average allelic expression. The escape threshold for dog was increased to exclude all bins within the dog putative pseudo-autosomal region.

### Modeling XCI ratios with the folded-normal distribution

Starting with a single parental allele, the sampled maternal allelic-expression of a heterozygous X-linked SNP can be modeled with a binomial distribution, dependent on the ratio of active maternal X-alleles in the sample and the read depth of the SNP.

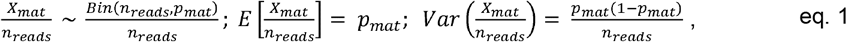

where *X*_*mat*_ is the number of maternal allelic reads, *n*_*reads*_ is the read depth of the SNP, and *p*_*mat*_ is the ratio of active maternal X-alleles. When aligned to a reference genome, the parental phasing information is lost and the allelic-expression of X-linked SNPs can instead be modeled with the folded-binomial model^59,60^. Since SNPs vary in read-depth, we use a folded-normal model as an approximation of the underlying mixture of depth-dependent folded-binomial distributions. The probability of allelic-expression under the folded-normal model is defined as:

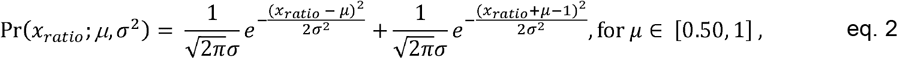

where *X*_*ratio*_ is the folded allelic-expression ratio of a SNP, *μ* is the folded XCI ratio of the sample, and *σ* is the standard deviation of the folded-normal distribution. We utilize a maximum-likelihood approach (negative log-likelihood minimization of eq. 2) to fit folded-normal distributions to the observed folded allelic-expression ratios of at least 10 filtered SNPs per sample, taking the *μ* parameter of the maximum-likelihood folded-normal distribution as the folded XCI ratio estimate of the sample.

### Modeling autosomal imbalances

The folded-normal model can also be applied to autosomal data to estimate allelic-imbalances. For each species, we extract chromosome-specific alignments from the .bam file for the two autosomes closest in size to the X-chromosome (Supp. Fig. 4). We employ the exact same processing pipeline and thresholds as used for the X-chromosome. Any sample that displayed an autosomal imbalance greater than or equal to a folded estimate of 0.60 (dotted lines in Supp. Fig. 4A) on either autosome was excluded from downstream analysis.

### Modeling population XCI variability with models of embryonic stochasticity

XCI is a binomial sampling event, where the number of cells choosing to inactivate the same X-allele follows a binomial distribution defined as:

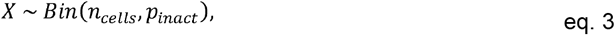

where *x* is the number of cells inactivating the same X-allele, *n*_*cells*_ is the number of cells present at the time of XCI, and *p*_*inact*_ is the probability of inactivation (0.50).

Embryonic XCI ratios can be modeled as:

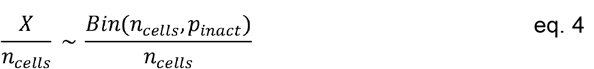

We estimate *n*_*cells*_ by fitting normal distributions to the unfolded population XCI ratio distributions of each species, as a continuous approximation for the underlying binomial distribution. The variance of the normal distribution is defined as:

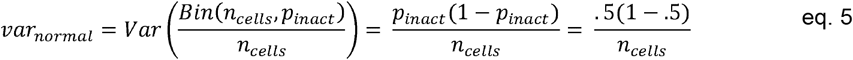

We model population XCI ratios as:

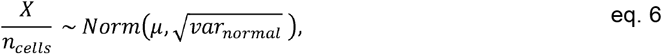

where *μ* = *p*_*inact*_ = 0.50 and *var*_*normal*_ is computed for *n*_*cells*_ ∈ [2, 200].

We identify the normal distribution with minimum sum-squared error between its CDF and the empirical population XCI ratio CDF, minimizing error over the tails of the distributions with percentiles <= 0.40 or >= 0.60 (Supp. Fig. 5). We compute 95% confidence intervals about the cell number estimate *n*_*cells*_ through bootstrap simulations. We sample with replacement from the empirical population XCI ratio distribution, matching the sample size of the original empirical population distribution, and fit a normal model to derive a bootstrap estimate of *n*_*cells*_. We repeat this for 2000 simulations to generate a bootstrapped distribution of *n*_*cells*_, from which we derive the 95% confidence intervals, defined as the interval where 2.5% of the bootstrapped distribution lies outside either end.

### Measuring sample X-linked heterozygosity

We compute sample heterozygosity as the ratio of SNPs detected in a sample (20 read minimum) to the total number of unique SNPs identified across all samples for a given species. We quantify associations between X-linked heterozygosity and XCI ratios as the spearman correlation coefficient between the sample X-linked heterozygosity ratio and the fitted mean and variance of the maximum-likelihood folded-normal distribution of the sample (Fig. 3B-C, Supp. Fig. 6). We only consider samples with at least 10 detected SNPs.

### Quantifying variant associations with extreme XCI ratios

We quantify the strength of XCI ratios as a predictor for the presence of a given variant through the AUROC metric. Given a ranked list of data (XCI ratios) and an indicator of true positives (samples with a given variant), the AUROC quantifies the probability a true positive is ranked above a true negative. An AUROC of 1 indicates all true positive samples were ranked above all true negative samples, demonstrating XCI ratios were a perfect predictor for the presence of that variant. An AUROC of 0.50 indicates random placement of true positives and negatives in the ranked list, demonstrating XCI ratios performed no better than random chance for predicting the presence of that variant. We compute the AUROC through the Mann-Whitney U-test, defined as:

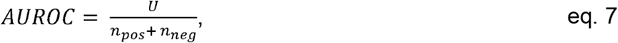

where *U* is the Mann-Whitney U-test test statistic, computed in R with wilcox.test(alternative = ‘two.sided’), *n*_*pos*_ is the number of true positive samples and *n*_*neg*_ is the number of true negative samples. We generate a null AUROC per variant by randomly shuffling the true positive and negative labels. The variant frequency is defined as the number of samples that carry a given variant over the total number of samples for a given species. The p-value for a given AUROC is the p-value associated with the Mann-Whitney U-test test statistic (*U*), where we determine significance as an FDR-corrected p-value <= 0.05. We perform FDR correction for all p-values computed for all variants across the 9 species through the Benjamini-Hochberg method, implemented in R via p.adjust(method = ‘BH’).

We estimate the power of each variant through bootstrap simulations. We randomly sample with replacement the XCI ratios of the true positive and true negative samples, those that either carry or do not carry a given variant. We match the sample size of the original true positive and negative labels. We compute a bootstrapped AUROC and p-value from the simulated data, repeating for 2000 simulations to compute a bootstrapped distribution of AUROCs. The AUROC power (Supp. Fig. 7B) is defined as the fraction of bootstrapped AUROCs that are significant, using a significance threshold of p-value <= 0.05. The AUROC effect size power (Supp. Fig. 7C) is defined as the fraction of bootstrapped AUROCs that are >= 0.75. We also report the variance of the bootstrapped AUROC distribution per variant in Supp. Fig. 7D. We exclude all variants classified as reference biased from Supp. Fig. 1, with the distributions of AUROCs for the reference biased and non-reference biased SNPs presented in Supp. Fig. 7E.

### Software

All analysis was performed in R^61^ v4.3.0. All plots were generated using ggplot2^62^ v3.4.2 functions. The phylogenetic tree in Fig. 2B was generated from TimeTree http://www.timetree.org/.

## Data and Code availability

All associated code can be found at https://github.com/gillislab/cross_mammal_xci. This includes the snakemake pipeline used for processing the non-human mammalian data as well as all R notebooks used for data analysis and figure generation.

## Author Contributions

J.G. conceived the project. J.M.W. and J.G. designed the experiments and wrote the manuscript. J.M.W. performed the experiments. J.H. and J.M.W performed data management and data processing.

## Acknowledgements

J.G., J.M.W., and J.H. were supported by NIH grants R01MH113005. We thank all members of the Gillis lab and particularly John Lee for assisting in some of the initial data downloading.

