## Supplemental Figures for "Population variability in X-chromosome inactivation across 9 mammalian species"

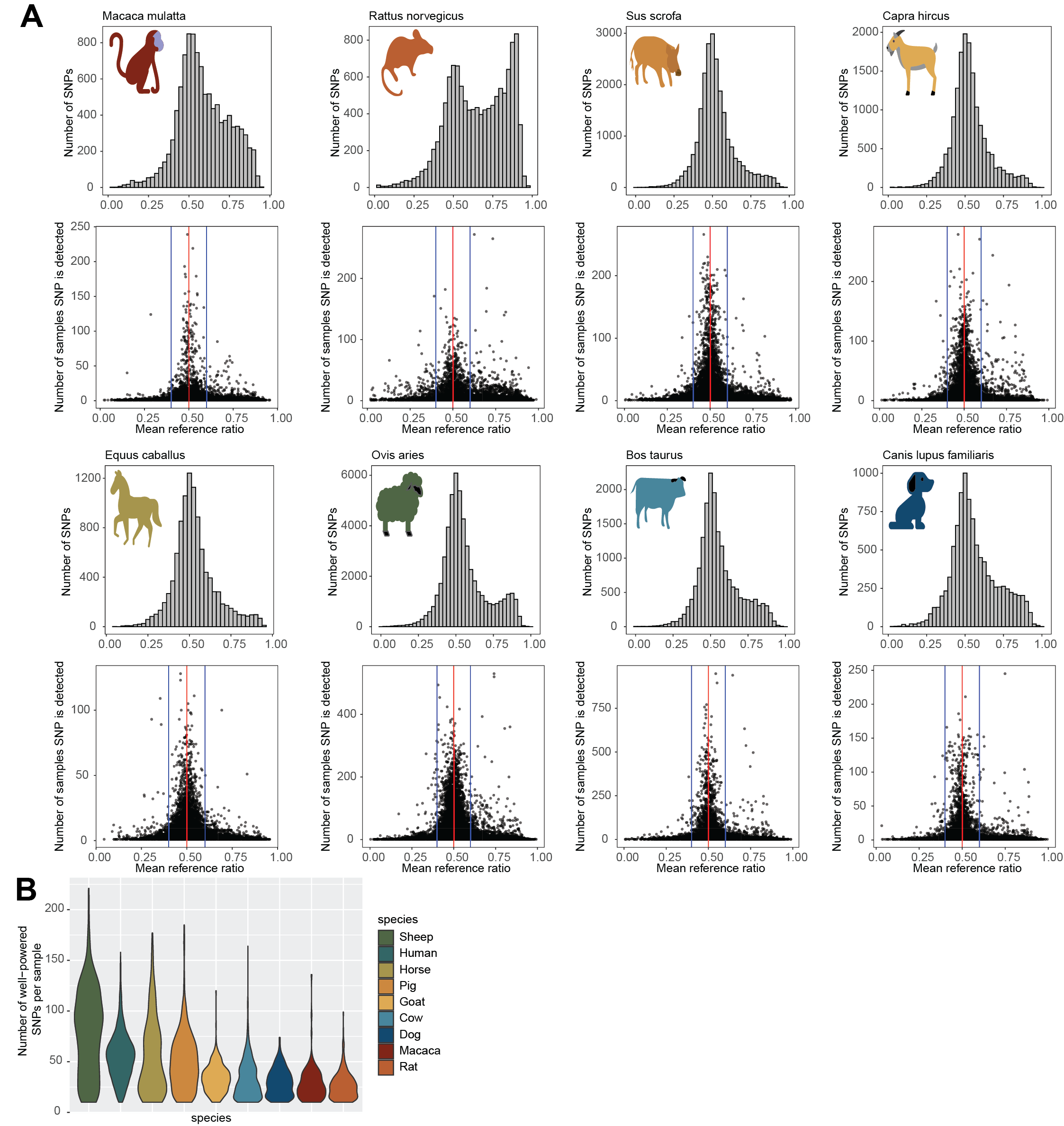

**Supplemental Figure 1: Reference bias varies across individual SNPs**

**A** Top histogram depicts the distribution of mean reference ratios for all detected SNPs in each species. The bottom scatter plot depicts the mean reference ratio against the sample size for each SNP. We exclude all SNPs from XCI ratio modeling whose mean reference ratio is < 0.40 or > 0.60 (blue lines), indicating consistent bias in allelic expression for either the alternate or reference allele.

**B** Violin plots depicting the distribution of the number of filtered SNPs (see methods) per sample for each species, where we require a minimum of 10 SNPs for XCI ratio modeling.

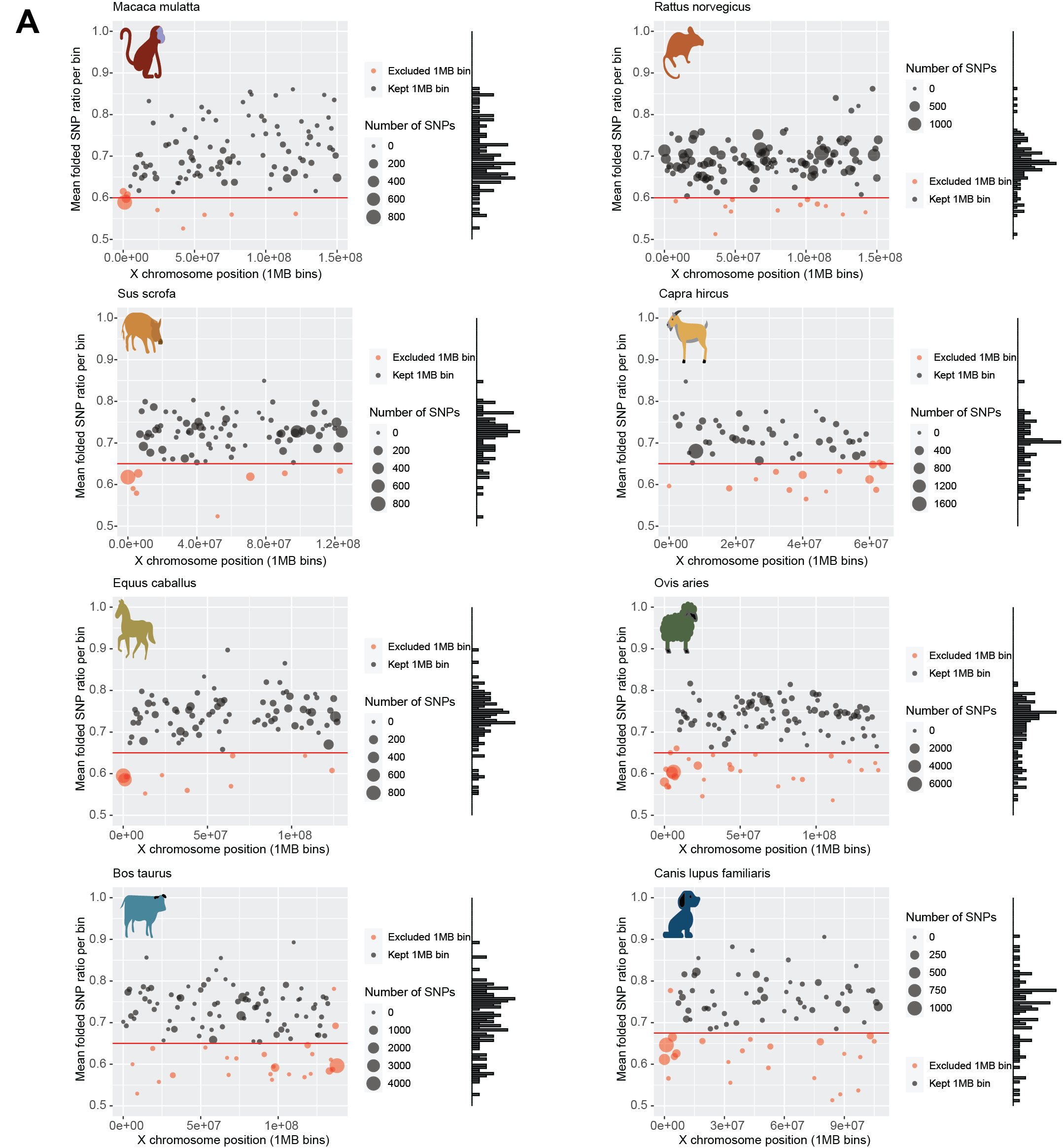

**Supplemental Figure 2: Escape from XCI is enriched in chromosomal ends**

**A** Scatter plots comparing the chromosomal location (1 mega-base bins) and the mean folded allelic expression ratio for all SNPs within each 1MB bin, derived from samples with skewed XCI (see methods). The marginal histogram depicts the distribution of mean folded allelic expression ratios per 1MB bin. The size of the data points corresponds to the number of SNPs in each 1MB bin. Red data points indicate 1MB bins that were excluded from analysis as probable escape regions, due to balanced allelic expression in samples with skewed XCI ratios. The chromosomal ends of all species, except rat, exhibit large clusters of SNPs with escape signal, likely pseudo-autosomal regions. The red lines depict the threshold of allelic-expression used to classify 1MB bins as escape or not.

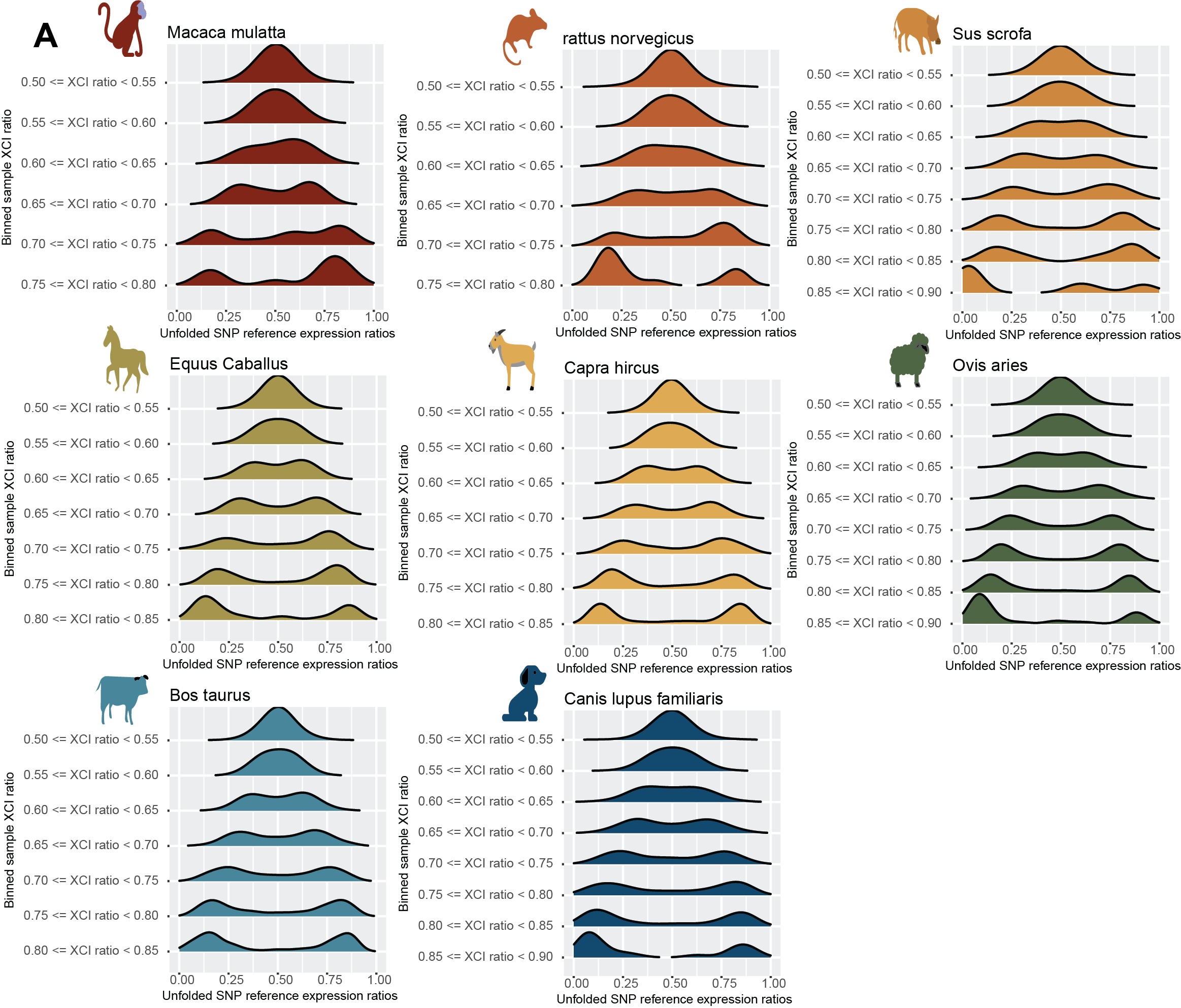

**Supplemental Figure 3: Reference allelic expression distributions exhibit bi-parental haplotype expression signatures expected of the X-chromosome**

**A** Density distributions of reference allelic expression ratios aggregated across samples binned by their estimated XCI ratio, ordered from balanced to more extreme XCI ratios (top to bottom). For samples with balanced XCI, the parental haplotypes cannot be distinguished, with clear separation of the parental haplotypes as the XCI ratio increases.

­

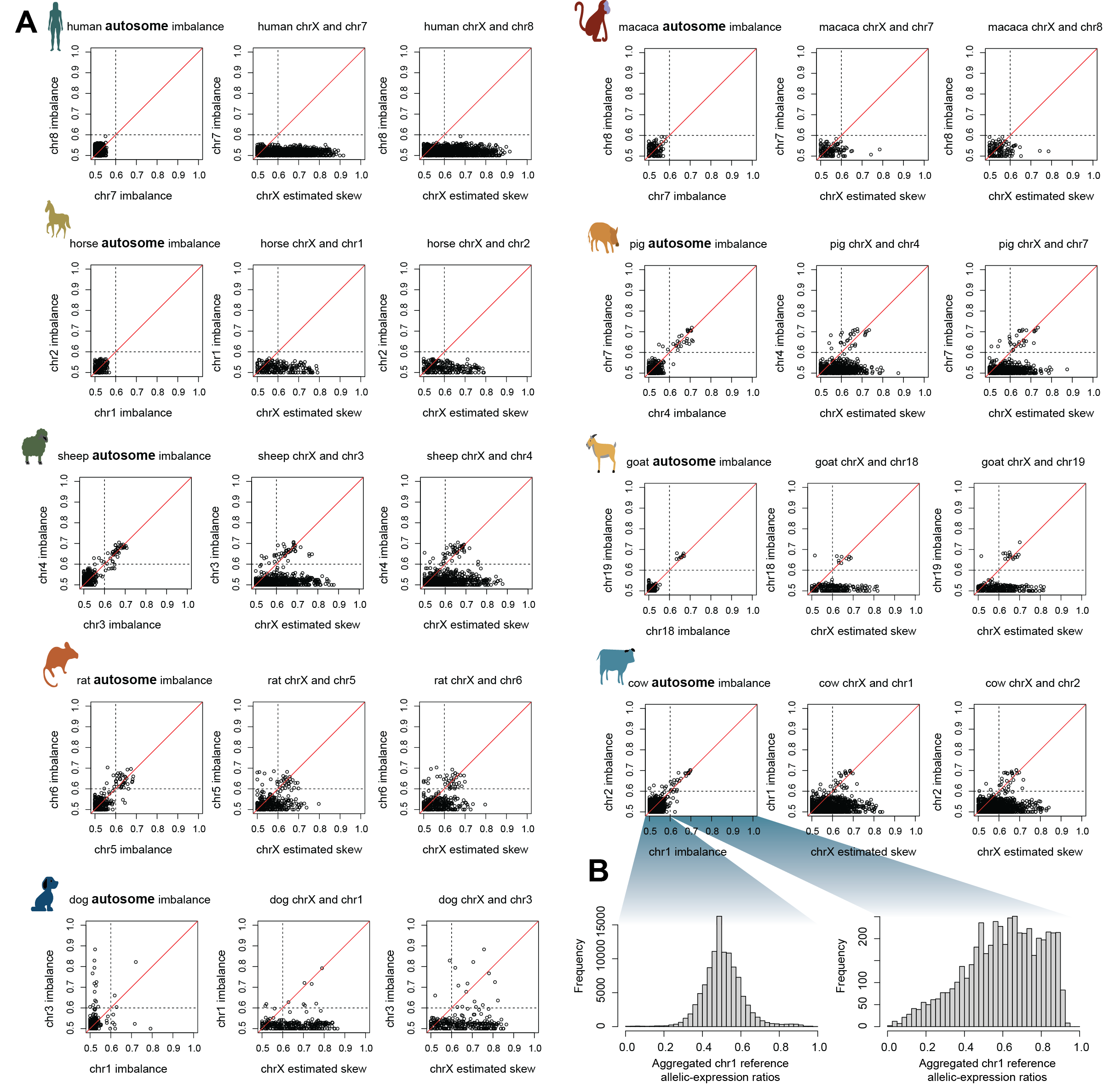

**Supplemental Figure 4: Comparing autosomal and X chromosome allelic imbalances**

**A** Each data point is a single individual sample. For each species, the first scatter plot compares the aggregated allelic imbalance of two autosomes (see methods). The following two scatter plots compare the aggregated allelic imbalance of an autosome and the X chromosome. Samples that exhibited imbalanced allelic expression on an autosome were excluded from analysis, using a threshold of an imbalance >= 0.60 (dotted lines).

**B** Histogram of the reference allelic-expression ratios of all chromosome 1 SNPs from the Cow samples with a chromosome 1 autosomal imbalance either < 0.60 (left) or >= 0.60 (right). The large autosomal imbalances can be attributed to extensive reference bias in allele-specific expression ratios. Cow results are representative of all other species.

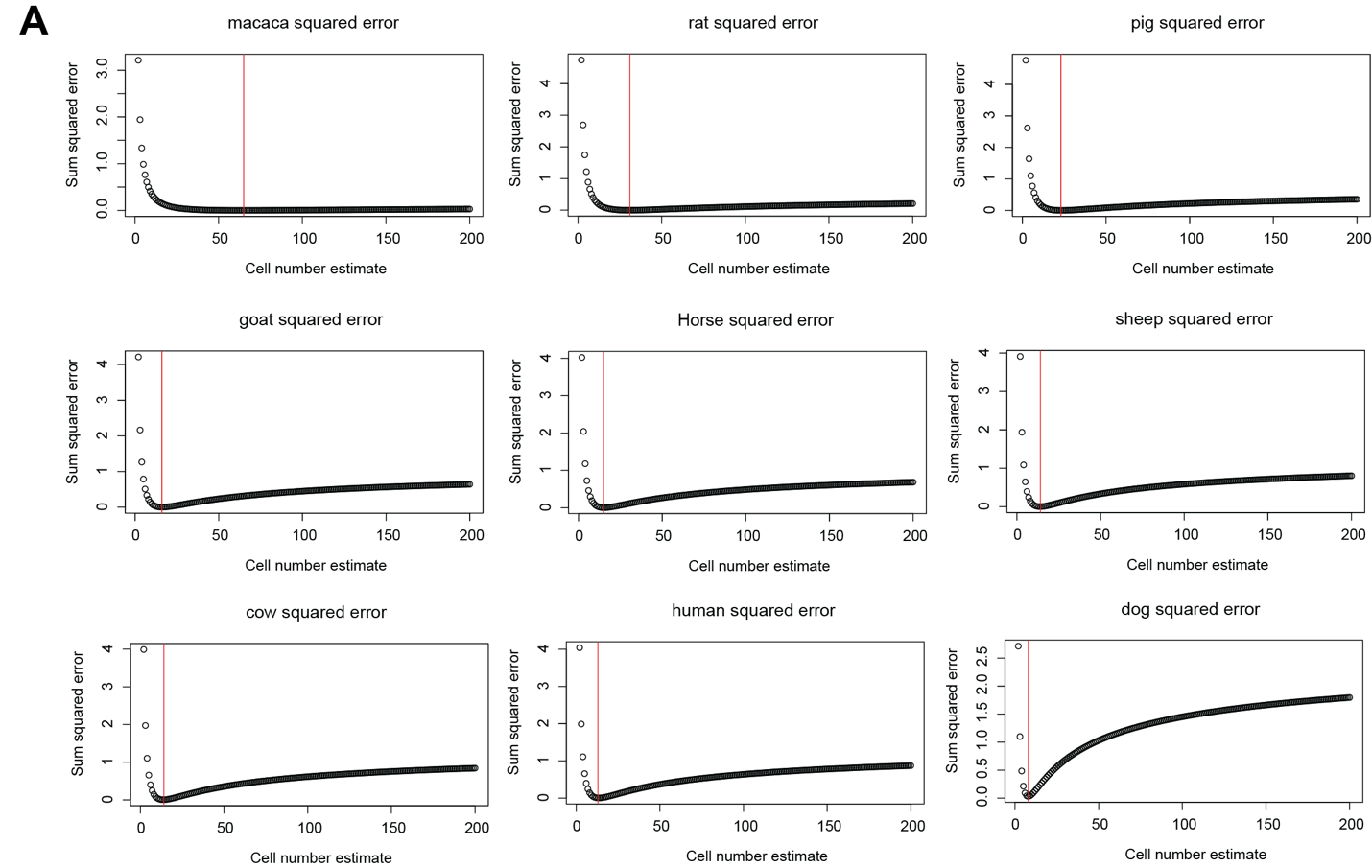

**Supplemental Figure 5: Estimating embryonic cell counts from population XCI ratio variance**

**A** Plots comparing the normal model associated with the estimated number of cells present during embryonic lineage specification (x-axis, see methods) to the sum of squared error between the percentiles of the tails of the empirical population XCI ratio distribution and the theoretical normal model (y-axis, see methods). The red lines depict the normal model with minimum error and the associated cell number estimate for each species.

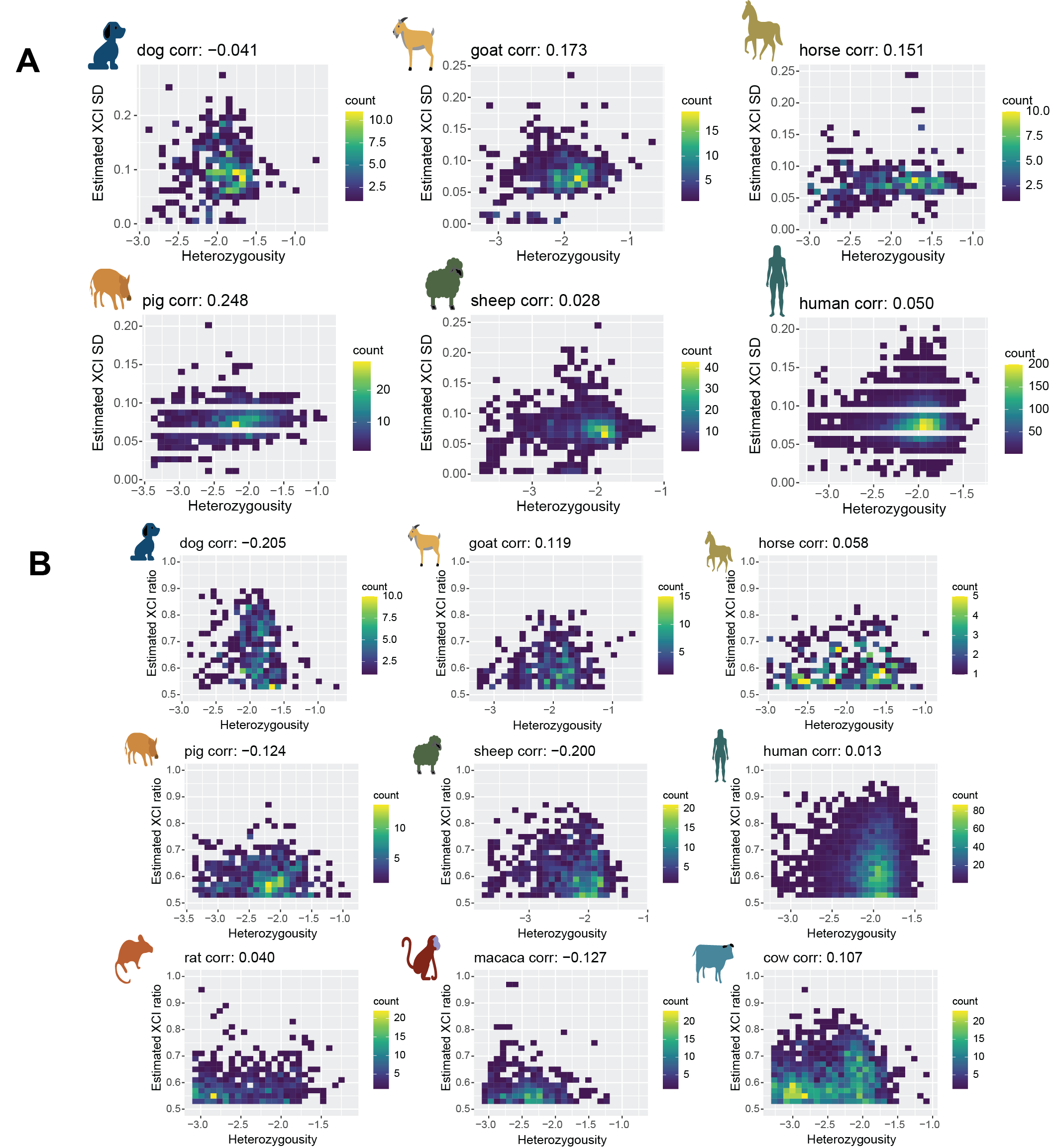

**Supplemental Figure 6: Species with no association between sample heterozygosity and variance in X-linked allelic expression**

**A** Binned scatter plots comparing the sample heterozygosity (log-10 of the number of SNPs per sample divided by the number of unique SNPs detected per species) to the estimated standard deviation (SD) in X-linked allelic expression (SD of the maximum-likelihood folded-normal distribution per sample). Spearman correlation coefficients are presented next to the species’ names. Color bars represent the number of datapoints per 2D bin.

**B** Binned scatter plots comparing the sample heterozygosity (log-10 of the number of SNPs per sample divided by the number of unique SNPs detected per species) to the estimated XCI ratio (mean of the maximum-likelihood folded-normal distribution per sample). Spearman correlation coefficients are presented next to the species’ names. Color bars represent the number of datapoints per 2D bin.

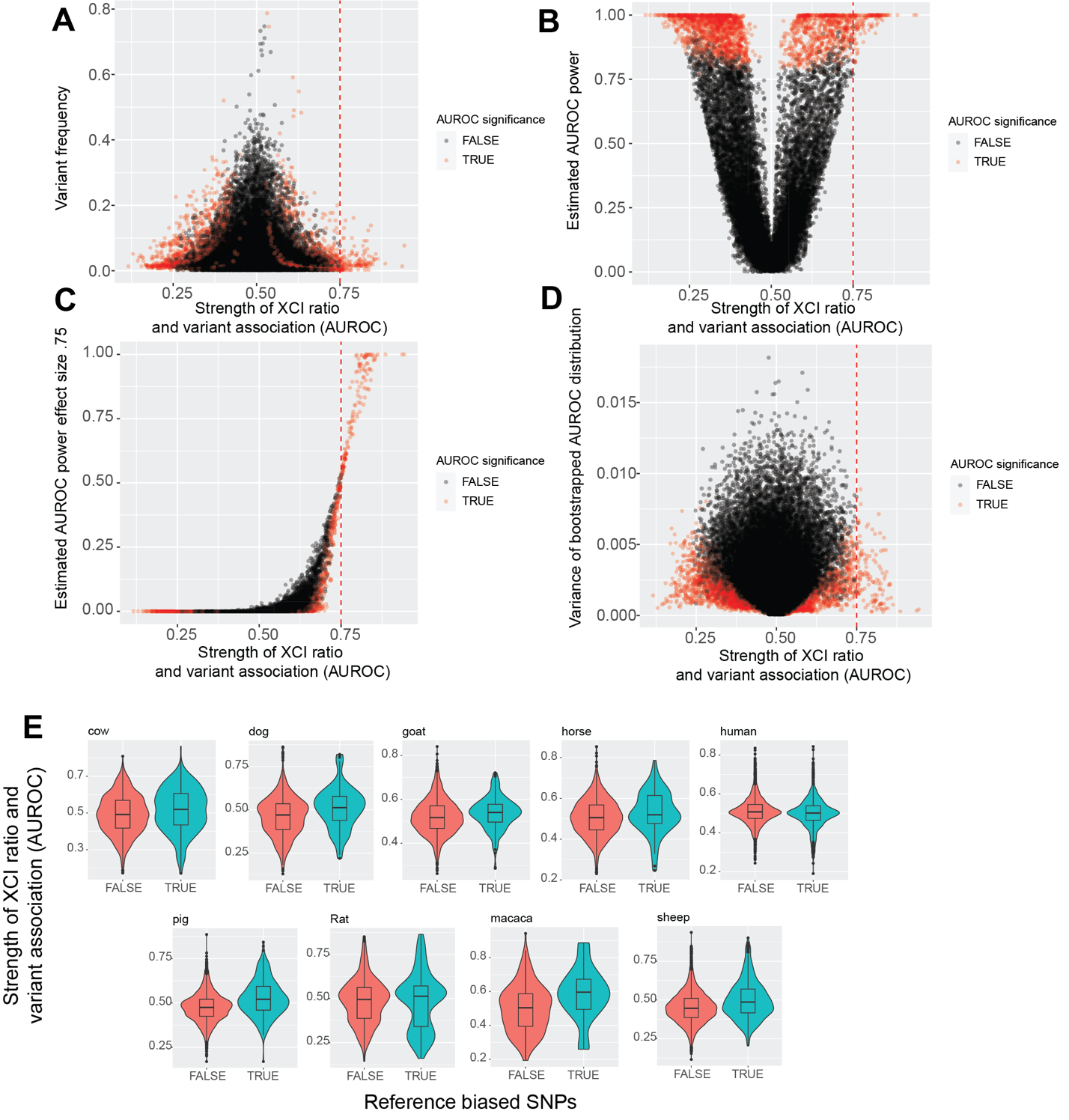

**Supplemental Figure 7: Low frequency variants are powered to detect significant associations with XCI ratios**

**A** Scatter plot comparing the initial AUROC against variant frequency for all variants across all species. Statistical significance of AUROCs is determined by an FDR-corrected p-value <= 0.05. The red dotted line in all 4 figure panels represents the AUROC threshold used to determine individual variants with a moderate association with XCI ratios.

**B** Scatter plot comparing the initial AUROC against the estimated power to detect a significant effect for each variant. Power was estimated through bootstrap simulations using a significance threshold of p-value <= 0.05, see methods.

**C** Scatter plot comparing the initial AUROC against the estimated power to detect an AUROC with effect size 0.75 or greater. Power was estimated through bootstrap simulations, see methods.

**D** Scatter plot comparing the initial AUROC against the variance of the bootstrapped distribution of AUROCs for each variant, see methods.

**E** Violin and boxplots depicting the distributions of AUROCs for the SNPs classified as either reference biased or not from the analysis in Supp. Fig. 1. Boxplots depict the quartiles of the distributions.
